# MOIRE: A software package for the estimation of allele frequencies and effective multiplicity of infection from polyallelic data

**DOI:** 10.1101/2023.10.03.560769

**Authors:** Maxwell Murphy, Bryan Greenhouse

**Affiliations:** Department of Biostatistics, School of Public Health, University of California, Berkeley, California, USA; EPPIcenter program, Division of HIV, ID and Global Medicine, University of California, San Francisco, California, USA

## Abstract

**Motivation:** Malaria parasite genetic data can provide insight into parasite phenotypes, evolution, and transmission. However, estimating key parameters such as allele frequencies, multiplicity of infection (MOI), and within-host relatedness from genetic data is challenging, particularly in the presence of multiple related coinfecting strains. Existing methods often rely on single nucleotide polymorphism (SNP) data and do not account for within-host relatedness.

**Results:** We present MOIRE (Multiplicity Of Infection and allele frequency REcovery), a Bayesian approach to estimate allele frequencies, MOI, and within-host relatedness from genetic data subject to experimental error. MOIRE accommodates both polyallelic and SNP data, making it applicable to diverse genotyping panels. We also introduce a novel metric, the effective MOI (eMOI), which integrates MOI and within-host relatedness, providing a robust and interpretable measure of genetic diversity. Extensive simulations and real-world data from a malaria study in Namibia demonstrate the superior performance of MOIRE over naive estimation methods, accurately estimating MOI up to 7 with moderate sized panels of diverse loci (e.g. microhaplotypes). MOIRE also revealed substantial heterogeneity in population mean MOI and mean relatedness across health districts in Namibia, suggesting detectable differences in transmission dynamics. Notably, eMOI emerges as a portable metric of within-host diversity, facilitating meaningful comparisons across settings when allele frequencies or genotyping panels differ. Compared to existing software, MOIRE enables more comprehensive insights into within-host diversity and population structure.

**Availability:** MOIRE is available as an R package at https://eppicenter.github.io/moire/.

**Supplementary information:** Supplementary data are available at *Bioinformatics* online.

## Introduction

Genetic data can be a powerful source of information for understanding malaria parasite phenotype and transmission dynamics, providing insight into population structure and connectivity, and thereby informing control and elimination efforts. However, analysis is complicated in malaria due to the presence of multiple coinfecting, genetically distinct strains. More specifically, genetically distinct strains may share the same alleles at genetic loci, rendering the actual number of strains contributing a particular allele unknown and making it difficult to estimate fundamental statistics such as population allele frequencies and multiplicity of infection (MOI). Standard methods to address this either naively estimate allele frequencies and MOI without considering the total number of strains contributing an allele (Roh *et al*., 2019; Tessema *et al*., 2019; Pringle *et al*., 2019), or completely ignore polyclonal samples during analysis. Naive estimation without accounting for strain count contribution results in biased estimates of allele frequencies and MOI, leading to meaningful systematic biases in summary statistics. For example, naive estimation of allele frequencies without consideration of strain composition from polyclonal samples results in a consistent overestimation of heterozygosity, leading to potentially faulty inference about population diversity. Additionally, naive estimation offers no principled way to address genotyping error beyond heuristics, further biasing estimates of diversity in ways that depend on choices made during initial interpretation of genotyping data. Alternatively, considering only monoclonal samples is potentially problematic, as this may require a substantial number of samples to be discarded when collected from regions where multiple infection is the rule rather than the exception. Further, the monoclonal subset of samples are fundamentally different from the larger population of interest, as they preclude the possibility of within-host relatedness between strains. This ignores a potentially important source of information about transmission dynamics, as within-host relatedness may be indicative of co-transmission or persistent local transmission (Wong *et al*., 2017; Nkhoma *et al*., 2020; Wong *et al*., 2018).

To address these issues and make full use of available data, Chang *et al*. (2017) developed a Bayesian approach (*THE REAL McCOIL*) to estimate allele frequencies and MOI in the context of polygenomic infections from single nucleotide polymorphism (SNP) based data. More recently, *coiaf* (Paschalidis *et al*., 2023) and *SNP-Slice* (Ju *et al*., 2023) have been developed to further improve computational efficiency and resolving power. Briefly, *coiaf* takes user provided allele frequencies and SNP read count data and applies an optimization procedure to estimate either discrete or continuous values for MOI. *SNP-Slice* also takes SNP read count data and uses a non-parametric Bayesian approach to simultaneously estimate phased strain identity and within-host strain composition. While Paschalidis et al. suggest within-host relatedness as a possible explanation for continuous values of MOI, and the method by Ju et al. may provide a way to interrogate within-host relatedness through phased strain composition, none of these methods directly consider or estimate within-host relatedness. Further, these methods are all tailored to SNP based data and are unable to accommodate more diverse polyallelic loci, such as microsatellites, which have been widely used in population genetic studies (Anderson *et al*., 2000; Tessema *et al*., 2019; Roh *et al*., 2019; Pringle *et al*., 2019). Other methods that infer within-host relatedness (Zhu *et al*., 2019), in contrast, rely on whole genome sequencing (WGS) data. WGS based approaches, however, frequently have poor sensitivity for detecting minority strains and low density infections (Tessema *et al*., 2022). In recent years, the declining cost of DNA sequencing and development of high throughput, high diversity, targeted sequencing panels have made polyallelic data even more attractive for genomic based studies of malaria (Tessema *et al*., 2022; LaVerriere *et al*., 2022; Kattenberg *et al*., 2023). Genetic analysis methods leveraging polyallelic loci have the potential for substantially increased resolving power over their SNP based counterparts, particularly in the context of related polyclonal infections in malaria (Taylor *et al*., 2019; Inna Gerlovina *et al*., 2022). Unfortunately, there are limited tools available to analyze these types of data.

We present here a new Bayesian approach, **M**ultiplicity **O**f **I**nfection and allele frequency **RE**covery from noisy polyallelic data (*MOIRE*), that, like *THE REAL McCOIL*, enables the estimation of allele frequencies and MOI from genomic data that are subject to experimental error. In addition, MOIRE estimates and accounts for within-host relatedness of parasites, a common occurrence due to the inbreeding of parasites serially co-transmitted by mosquitoes (Nkhoma *et al*., 2020, 2012). Critically, MOIRE takes as input genetic data of arbitrary diversity, allowing for estimation of allele frequencies, MOI, and within-host relatedness from polyallelic as well as biallelic data. MOIRE is able to fully utilize polyallelic data, yielding joint estimates of allele frequencies, sample specific MOIs and within-host relatedness along with probabilistic measures of uncertainty. We demonstrate through simulations and applications to empirical data the ability of MOIRE to leverage a variety of polyallelic markers. Polyallelic markers can greatly improve jointly estimating sample MOI, within-host relatedness, and population allele frequencies, resulting in reduced bias and increased power for understanding population dynamics from genetic data. We also introduce a new metric of diversity, the effective MOI (eMOI), a continuous value that combines estimates of the true MOI and the degree of within-host relatedness in a single sample, providing an interpretable quantity that is comparable across genotyping panels and transmission settings. We contrast this with the within-host infection fixation index, *F*_*W S*_, a frequently used metric of within-host diversity and signal of inbreeding and population sub-structure (Manske *et al*., 2012; Auburn *et al*., 2012), and demonstrate the inherent shortcomings of *F*_*W S*_ as a non-portable metric.

## Methods

### A model of infection and observation

Consider observed genetic data *X* = (*X*_1_, …, *X*_*n*_) from *n* samples indexed by *i*, where each *X*_*i*_ is a collection of vectors indexed by *l* of possibly differing length, representing the varying number of alleles possible at each locus, e.g. polyallelic loci. Each vector is binary, with 1 representing the allele was observed or 0 representing the allele went unobserved at locus *l* for sample *i*. From this data, we wish to estimate MOI for each individual (***µ*** = [*µ*_1_, …, *µ*_*n*_]), within host relatedness (***r*** = [*r*_1_, …, *r*_*n*_]), defined as the average proportion of the genome that is identical by descent across all strains, individual specific genotyping error rates 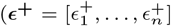 and 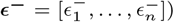, and population allele frequencies at each locus (***π*** = [*π*_1_, …, *π*_*l*_]). Similar to Chang *et al*. (2017), we applied a Bayesian approach and looked to estimate the posterior distribution of ***µ, r, ϵ***^**+**^, ***ϵ***^−^ and ***π*** as

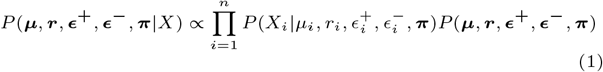

where we assumed independence between samples.

Given that the observed genetic data are experimentally derived, they are subject to some rate of false positives where an allele is erroneously called as present, and false negatives where an allele is erroneously called as absent. To address this issue, we augmented our model with a latent true genetic state *Y*, reflecting the true presence or absence of alleles at each locus for each individual. Augmenting our model with this latent state allowed us to incorporate and model the uncertainty around measurement of genetic data separately from the uncertainty around the true genetic state, as expressed in the following factorization:

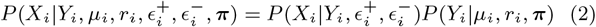

We assumed a prior in which the MOI of each individual was independent of the MOI of other individuals, relatedness was independent across individuals, error rates were independent across individuals, and allele frequencies were independent across loci and without linkage disequilibrium, yielding the following factorization:

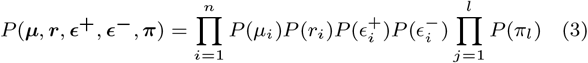

Details of the likelihood and prior distributions are provided in the supplementary material (section 1 and section 2), as well as practical considerations when using MOIRE with real world data (section 3).

### Effective MOI

By estimating MOI and within-host relatedness, we can estimate a continuous metric of genetic diversity within a host, the effective MOI (eMOI), which we define as:

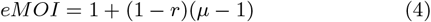

One interpretation of the effective MOI is the expected number of distinct alleles at a locus with infinite diversity, i.e. a locus where heterozygosity is 1 (see supplementary section 6 for a formal derivation). In the case of no within-host relatedness, this is simply the MOI. However, when there is within-host relatedness, the effective MOI is the MOI weighted by the probability that a given strain is unrelated to all other strains within the host, and ranges from 1 to *µ*. This value better reflects the true genetic diversity within a host than the MOI alone, and allows for comparison and differentiation of genetic diversity across hosts with the same MOIs. We also note that eMOI is likely to be more identifiable than MOI or within-host relatedness alone because it is a one-dimensional combination of the two estimated parameters with synergistic properties around precision. As MOI increases, precision around estimates of within-host relatedness also increases as there are more observations available to inform within-host relatedness. As MOI decreases, precision around the estimate of within-host relatedness decreases, however the contribution to the estimate of eMOI from within-host relatedness also decreases, and thus the overall precision of eMOI is maintained.

### Inference and Implementation

We fit our model to observed genetic data using a Markov Chain Monte Carlo (MCMC) approach using the Metropolis-Hastings algorithm with a variety of update kernels. Details of sampling and implementation are described in the supplementary material (section 5). MOIRE is implemented as an R package and is available with tutorials and usage guidance at https://eppicenter.github.io/moire/. All sampling procedures were implemented using Rcpp (Eddelbuettel and Francois, 2011) for efficiency. Substantial effort was placed on ease of use and limiting the amount of tuning required by the user by leveraging adaptive sampling methods. We provide weak default priors for all parameters and recommend that users only modify priors if they have strong prior knowledge about the parameters, such as experimentally derived estimates of false positive and false negative rates using samples with known parasite compositions and densities. All analysis conducted in this paper was done using MOIRE with default priors and settings, using 40 parallel tempered chains for 5000 burn-in steps, followed by 10,000 samples which were thinned to 1000 total samples.

## Results

### Estimation of multiplicity of infection, within-host relatedness, and allele frequencies

We simulated collections of 100 samples under varied combinations of population mean MOI, average within-host relatedness, false positive and false negative rates, and different genotyping panels (details of our simulation procedure may be found in the supplement section 4). Individual MOIs were drawn from zero truncated Poisson (ZTP) distributions with rate parameters 1, 3, and 5, resulting in mean MOIs of 1.58, 3.16, and 5.03 respectively. Within-host relatedness was simulated from settings with low, moderate, and high relatedness. False positive and false negative rates were varied from 0 to 0.1. We first simulated synthetic genomic loci with prespecified diversity: 100 SNPs, 30 loci with 5 alleles (moderate diversity), 30 loci with 10 alleles (high diversity), and 30 loci with 20 alleles (very high diversity) with frequencies drawn from the uniform Dirichlet distribution. We also assessed potential real world performance of MOIRE by simulating data for 5 currently used genotyping panels from 12 regional populations characterized by the MalariaGEN Pf7 dataset (Abdel Hamid *et al*., 2023) as described in the supplementary material (section 7, Supplementary Figure 4). Genetic loci were selected according to a 24 SNP panel (Daniels *et al*., 2008), a 101 SNP panel (Chang *et al*., 2019), and 3 recently developed amplicon sequencing panels consisting of 128 (LaVerriere *et al*., 2022), 165 (Aranda-Diaz and Neubauer Vickers, 2022), and 233 (Kattenberg *et al*., 2023) diverse microhaplotypes respectively. Like the fully synthetic simulations, these simulations were varied over a range of MOI and within-host relatedness, however error rates were fixed at moderate false positive and false negative rates of .01 and .1 respectively for the purposes of computational feasibility due to the extensive number of simulations required. We chose these levels as we believe they are reflective of the most likely situation of higher levels of false negatives and relatively low rates of false positives from a typical bioinformatics pipeline. We then ran MOIRE and calculated summary statistics of interest on the sampled posterior distributions.

We estimated allele frequencies, heterozygosity, MOI, within-host relatedness, and eMOI using the mean or median of the posterior distribution output by MOIRE. It should be noted that within-host relatedness is only defined for polyclonal infections, so the posterior distribution of within-host relatedness is conditional on the MOI being greater than 1. We contrasted these with naive estimates of allele frequency and MOI by assuming that an observed allele was contributed by a single strain, and estimated MOI as equal to the second-highest number of alleles observed across loci. We calculated ground truth allele frequencies using the true number of strains contributing each allele.

Under moderate false positive and false negative rates of 0.01 and 0.1 respectively, MOIRE accurately recovered parameters of interest across a range of genotyping panels, population MOI, and within-host relatedness (Figure 1, Table 1). Allele frequencies estimated by MOIRE were unbiased across genotyping panels (Figure 1B), leading to unbiased estimates of heterozygosity (Figure 1C). Naive estimation exhibited substantial bias that varied with respect to the true allele frequency. Rare alleles tended to be overestimated and common alleles underestimated, leading to inflated estimates of heterozygosity.

**Table 1.** Mean absolute deviation (MAD) of estimates of MOI, heterozygosity, within-host relatedness, and eMOI across simulations using synthetic (top) and real-world (bottom) genotyping panels. The MAD of estimates of MOI were calculated by taking the mean of the MAD for each stratum of true MOI between 1 and 10. MOI Within-host relatedness accuracy is only considered for samples with a true MOI *>* 1. Coverage rates of 95% credible intervals are shown in parentheses for estimates by MOIRE.

| Panel | Source | Heterozygosity |  | Allele Freqs. |  | MOI |  | Relatedness | eMOI |
| --- | --- | --- | --- | --- | --- | --- | --- | --- | --- |
|  |  | MOIRE | Naive | MOIRE | Naive | MOIRE | Naive | MOIRE | MOIRE |
| 100 SNP | Synthetic | 0.01 (.95) | 0.05 | 0.02 (.95) | 0.06 | 1.72 (.85) | 3.61 | 0.20 (.70) | 0.37 (.77) |
| Moderate Div. | Synthetic | 0.01 (.99) | 0.04 | 0.01 (.96) | 0.02 | 1.55 (.88) | 2.53 | 0.14 (.77) | 0.17 (.91) |
| High Div. | Synthetic | 0.01 (.99) | 0.02 | 0.01 (.91) | 0.01 | 1.29 (.86) | 1.87 | 0.11 (.75) | 0.12 (.89) |
| Very High Div. | Synthetic | 0.02 (.60) | 0.01 | 0.01 (.82) | 0.01 | 1.02 (.86) | 1.28 | 0.10 (.77) | 0.10 (.85) |
| 24 SNP | Daniels et al. (2008) | 0.01 (.90) | 0.04 | 0.02 (.90) | 0.05 | 1.95 (.81) | 3.66 | 0.21 (.75) | 0.45 (.86) |
| 101 SNP | Chang et al. (2019) | 0.01 (.90) | 0.04 | 0.02 (.95) | 0.05 | 1.77 (.85) | 3.62 | 0.20 (.71) | 0.36 (.79) |
| AMPLseq | LaVerriere et al. (2022) | 0.01 (.97) | 0.05 | 0.01 (.94) | 0.02 | 1.32 (.88) | 1.86 | 0.12 (.71) | 0.14 (.88) |
| MaD<sup>4</sup>HatTeR | Aranda-Diaz and Neubauer Vickers (2022) | 0.01 (.98) | 0.05 | 0.01 (.95) | 0.03 | 1.24 (.90) | 1.88 | 0.13 (.68) | 0.12 (.88) |
| AmpliSeq | Kattenberg et al. (2023) | 0.02 (.94) | 0.06 | 0.01 (.93) | 0.02 | 1.34 (.83) | 1.56 | 0.12 (.67) | 0.12 (.88) |

**Fig. 1:**
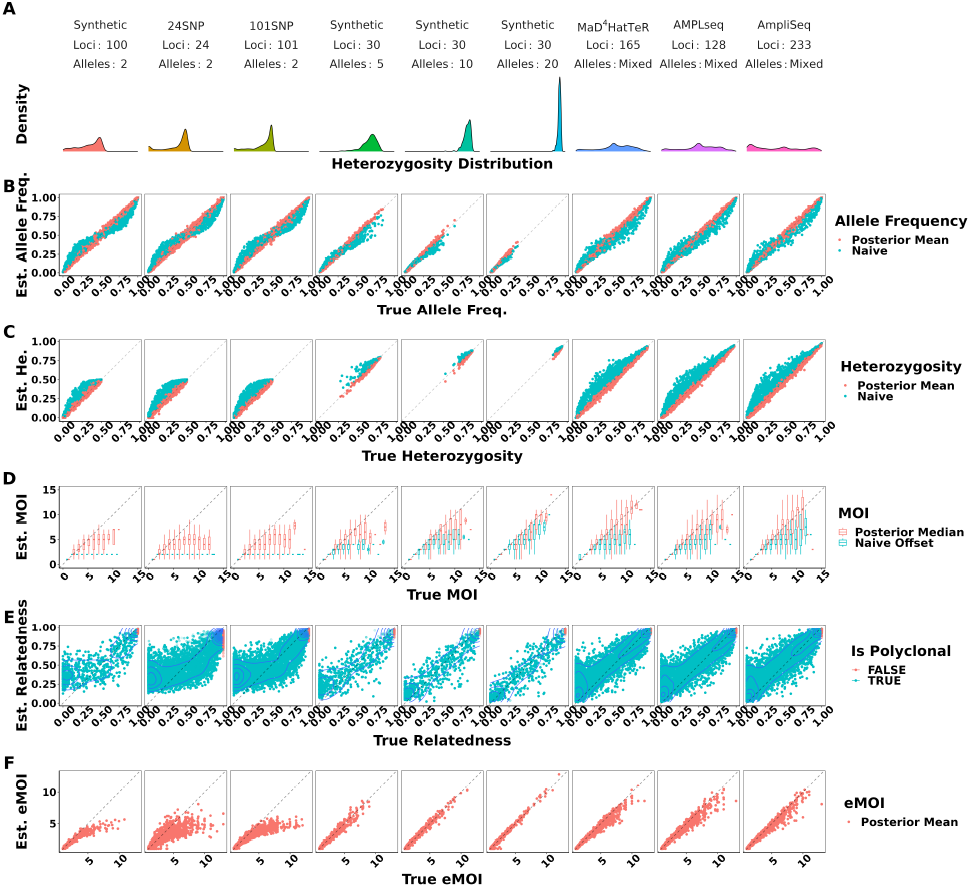
True vs. estimated values of parameters across panels of varying genetic diversity. Panel A summarizes the distribution of heterozygosity across each panel used. Each symbol represents the estimated value of the parameter for a single simulated dataset, with the true value of the parameter on the x-axis and the estimated value on the y-axis. Simulations were pooled across mean MOIs and levels of relatedness. False positive and false negatives rates were fixed to 0.01 and 0.1 respectively. Opacity was set to accommodate overplotting, except in the case of within-host relatedness where opacity reflects the estimated probability that a sample is polyclonal, calculated as the posterior probability of the sample MOI being greater than 1, as individual within-host relatedness is only defined for samples with MOI greater than 1. MOIRE accurately recovered parameters of interest with increasing accuracy as panel diversity increased, while naive estimation exhibited substantial bias where such estimators exist.

MOI was also well estimated by MOIRE, with accuracy increasing substantially in the presence of more diverse loci (Figure 1D). In the context of SNPs, MOIRE recovered MOI accurately up to approximately 4 strains, and then began to exhibit limited ability to resolve. More diverse panels enabled greatly improved resolving power, allowing for the accurate recovery of MOI up to approximately 7 strains. Naive estimation substantially underestimated MOI in comparison, due in part to the limited capacity of low diversity loci to discriminate MOI, as well as the presence of related strains that deflate the observed number of distinct alleles. This bias was particularly prominent for low diversity markers such as SNPs which can only resolve up to 2 strains.

MOIRE was generally able to recover within-host relatedness, particularly for moderate and high diversity markers in the context of high relatedness (Figure 1E). SNP based panels had difficulty resolving individual level within-host relatedness and were sensitive to the uniform prior. It should be noted that in the circumstance that a monoclonal infection has an inferred MOI greater than 1, MOIRE will likely classify these infections with very high relatedness (Figure 1E). This is due to the presence of false positives that MOIRE will sometimes infer as an infection consisting of highly related strains rather than being explained by observation error. Therefore, within-host relatedness should be interpreted in the context of the probability of the infection being polyclonal. A more robust metric is eMOI, since it is a metric of diversity that integrates MOI and within-host relatedness.

MOIRE recovered eMOI with high accuracy under all conditions using polyallelic panels (Figure 1F). SNP panels exhibited a larger degree of bias at higher eMOI, but still performed relatively well for eMOI of up to 4. This demonstrates that while identifiability of MOI or within-host relatedness may be challenging in some situations, eMOI is a reliably identifiable quantity when estimated using highly polymorphic markers.

All simulations were also conducted without any relatedness present. MOIRE was still able to accurately recover allele frequencies, heterozygosity, and MOI, indicating that minimal bias or uncertainty are introduced by attempting to estimate relatedness (Supplementary Figure 1).

These patterns held across the range of false positive and false negative rates simulated with the fully synthetic simulations.

Allele frequencies and heterozygosity remained well estimated by MOIRE across settings, however bias was elevated for individual level estimates of MOI, within-host relatedness, and eMOI when false positive rates were increased and panel diversity was low. Increased false negative rates did not result in any additional bias within the range of tested values (Supplementary Figure 2).

### Population inference

MOIRE is a probabilistic approach providing a full posterior distribution over model parameters, allowing estimation of credible intervals for model parameters as well as functions thereof. While sample level parameters estimated by the model are useful, it may also be useful to estimate population level summary statistics for reporting and comparison purposes. We thus calculated the posterior distribution of population level summaries of interest, such as mean MOI, mean within-host relatedness, and mean eMOI. We note that mean within-host relatedness is defined only for samples with MOI greater than 1, therefore the posterior distribution of mean within-host relatedness was calculated across samples with MOI greater than 1 at each iteration of the MCMC algorithm. MOIRE accurately estimated these quantities across a range of conditions (Supplementary Figure 3), with the best performance seen for polyallelic data.

Population mean MOI was accurately estimated across all panels, with improved precision at lower levels (Supplementary Figure 3A, Table 1). Credible interval (CI) coverage in general was poor, likely due to the challenge of identifiability in conjunction with within-host relatedness. SNP panels were largely unable to resolve population level mean within-host relatedness and exhibited poor CI coverage and substantial sensitivity to the uniform prior specification due to the low relative information contained in these markers. Polyallelic panels in contrast had improved precision as more diverse panels were used, although CI coverage was also poor due to persistent sensitivity to the uniform prior as indicated by slightly overestimating within-host relatedness below .5 and underestimating within-host relatedness above .5.

Population mean eMOI was remarkably accurate for low and medium mean MOI when using SNP based panels, with bias only becoming apparent at higher mean MOI (Supplementary Figure 3C, Table 1). Polyallelic panels had substantially improved precision across a wide range of values, further demonstrating that while population mean within-host relatedness or mean MOI may be challenging to identify, mean eMOI remains a highly identifiable quantity when genetic markers with sufficient diversity are used.

### Metric stability across genetic backgrounds

Population metrics of genetic diversity enable researchers to make comparisons across space and time, and to answer questions relating to differences in transmission dynamics. In order for a metric to be useful for these purposes, it must be sensitive to changes in transmission dynamics while remaining insensitive to other factors that vary and may confound interpretation, such as the genotyping panel used, or the local allele frequencies for a given panel. For example, if we were to compare two populations that exhibit the same transmission dynamics, we would want the metric to be the same, uninfluenced by differing population allele frequencies. It would be even better if the metric is insensitive to the genotyping panel used, allowing for comparisons across studies that are independent of the technology utilized.

To explore the performance of eMOI across varying transmission settings, we simulated 100 samples with MOI drawn from a ZTP distribution with either *λ* = 1 or *λ* = 3. For each sample, we then simulated either low or high within-host relatedness. For each individual level simulation, we then observed simulated genetics parameterized by each of the 12 regional populations previously described using the 5 genotyping panels, followed by the previously described observation process with false positive and false negative rates of .01 and .1 respectively. We then fit MOIRE on each simulation independently.

For each simulation, we calculated mean eMOI, mean naive MOI, and the within-host infection fixation index (*F*_*W S*_) (Roh *et al*., 2019; Manske *et al*., 2012), a frequently used metric of within-host diversity that relates genetic diversity of the individual infection to diversity of the parasite population. Mean MOI was calculated using the second-highest number of observed alleles, and *F*_*W S*_ used the observed genetics, assuming all alleles were equifrequent within hosts, and naive estimates of allele frequencies to estimate heterozygosity. For these metrics to be most useful in characterizing transmission dynamics, they should be the same for all simulations with the same degree of within-host relatedness and mean MOI, no matter the panel used nor the genetic background of the population. We found that mean eMOI was stable across all genetic backgrounds using microhaplotype based panels, yielding accurate estimates of mean eMOI despite substantial variability in local diversity of alleles, as shown by heterozygosity, and differing genomic loci (Figure 2A). Interestingly, while the SNP panels exhibited reduced precision and downward bias as expected, they were consistently biased with respect to the true eMOI, even across different panels. This suggests that SNP panels, while limited in resolving power, may still have utility in estimating relative ordering of eMOI. These results also demonstrate that eMOI may be readily used and compared across transmission settings and is relatively insensitive to other factors such as heterozygosity that may vary across settings. In contrast, mean naive MOI and *F*_*W S*_ were sensitive to genetic background and genotyping panel in confounded ways. Mean naive MOI, only useful with polyallelic markers, exhibits an inherent upward bias as mean heterozygosity increases that is most severe at higher mean MOI. This bias also varied with the genotyping panel used, making it difficult to interpret and compare across settings (Figure 2B). *F*_*W S*_ is also sensitive to genetic background and panel used, exhibiting an upward trend as heterozygosity increases and a bias that varies across panels. This is inherent to the construction of the metric, as it is coupled to an estimate of the true heterozygosity of genetic loci being used (Figure 2C). This simulation demonstrates limitations in the utility of *F*_*W S*_ as a metric of within-host diversity for a population as it is inherently uncomparable across settings due to its high sensitivity to varying genetic background and genotyping panel used. Mean eMOI, in contrast, is a stable metric of genetic diversity that is insensitive to genetic background and genotyping panel, and is thus readily comparable across settings.

**Fig. 2:**
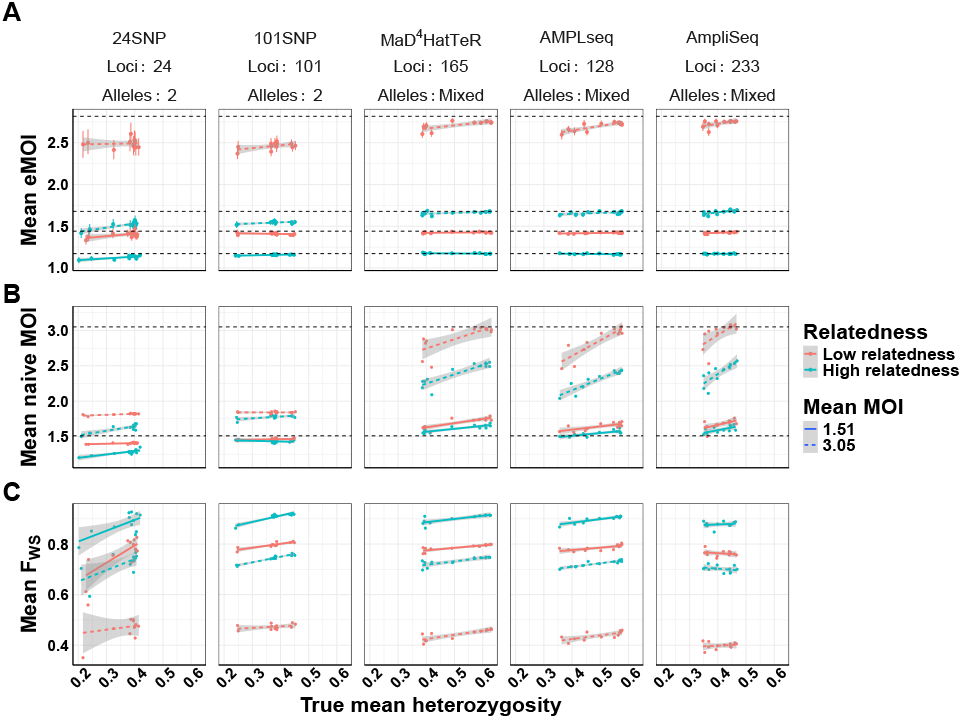
Comparison of mean eMOI to other summary measures of diversity across varying levels of within-host relatedness. For each level of relatedness (low and high), we simulated 100 infections with a mean MOI of 1.51 and 3.16, for a total of 400 infections across 4 conditions. Keeping the MOI and relatedness fixed for each sample, we varied the genetic diversity of the panel used to genotype each sample. We then calculated the mean eMOI from MOIRE, mean MOI using the naive estimator, and mean *F*_*W S*_ using a naive estimate of allele frequencies for each simulation to assess the sensitivity of each metric to varying the genetic diversity of the panel. True mean eMOI and mean MOI are fixed values within levels of within-host relatedness and are annotated by dashed lines. Mean *F*_*W S*_ is not fixed within levels of within-host relatedness and MOI because it is a function of the genetic diversity of the panel.

### Application to a study in Northern Namibia

We next used MOIRE to reanalyze data from a previously conducted study carried out in northeastern Namibia consisting of 2585 samples from 29 health facilities across 4 health districts genotyped at 26 microsatellite loci (Tessema *et al*., 2019). We ran MOIRE across samples collected from each of the 4 health districts independently. Running MOIRE in this way implies that we are assuming that all samples from each health district come from a shared population with the same allele frequencies. We then calculated summary statistics of interest on the sampled posterior distributions.

We compared our results to the naive estimation conducted in the original study and found that overall relative ordering of mean MOI was maintained, with Andara and Rundu exhibiting the highest MOI, Zambezi the lowest, and Nyangana in between, consistent with contemporary estimates of transmission intensity (Tessema *et al*., 2019). However, similar to our simulations, naive estimation substantially underestimated mean MOI across health districts compared to MOIRE (Figure 3A and C). Individual within-host relatedness was estimated to be very high across sites (IQR: .61-.91) with no differences between sites (Figure 3B). This suggests substantial inbreeding which may be indicative of persistent local transmission, consistent with the original findings by Tessema *et al*. (2019) We also found that heterozygosity across loci estimated by MOIRE was generally lower (IQR: .55 - .85), consistent with the previously described simulations demonstrating that naive estimation overestimates heterozygosity, and that previously detected statistically significant differences in heterozygosity between the Zambezi region and the other three regions may have been an artifact of biased estimation (Figure 3D).

**Fig. 3:**
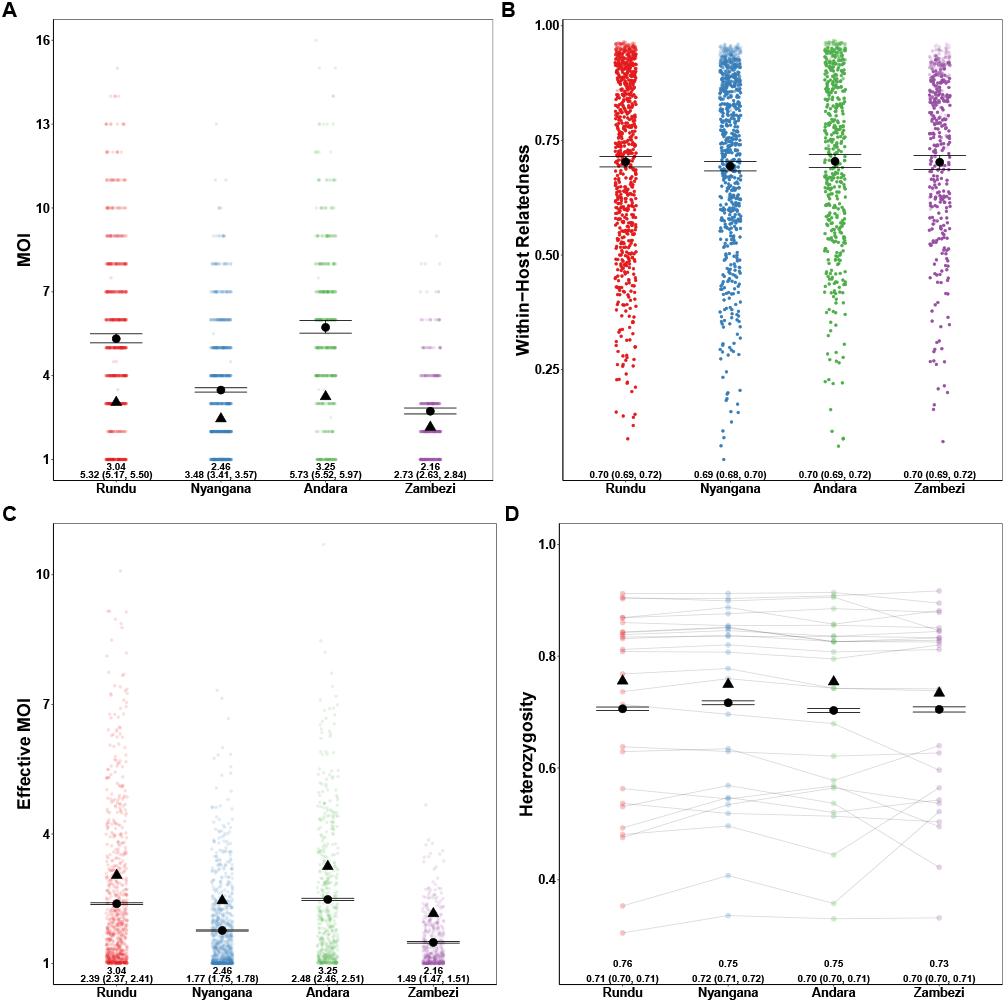
Estimated MOI, relatedness, eMOI and heterozygosity in Northern Namibia. MOIRE was run on data from 2585 samples from 29 clinics genotyped at 26 microsatellite loci, subset across four health districts. Each point represents the posterior mean or median for each sample or locus level parameter. The black circle represents the population mean with 95% credible interval for each health district and the black triangle indicates the naive estimate where applicable. In the case of eMOI (C), the naive estimate is simply the MOI. Opacity was used to accommodate overplotting in A, C and D, however opacity in B is reflective of the posterior probability of a particular sample being polyclonal to emphasize that an observation’s contribution to the posterior distribution of mean within-host relatedness is weighted by its probability of being polyclonal. This is due to the fact that mean within-host relatedness is only defined for samples with MOI greater than 1, and thus the posterior distribution of within-host relatedness was calculated by taking the mean within-host relatedness across samples with MOI greater than 1 at each iteration of the MCMC algorithm. Therefore, the opacity of each point in B is reflective of the contribution of that sample to the posterior distribution of mean within-host relatedness.

We also ran MOIRE independently across each of the 29 health facilities, excluding 2 health facilities from the Zambezi region due to low total number of samples (n = 9 in each). Stratifying by health facility revealed substantial heterogeneity in mean MOI, within-host relatedness, and consequently eMOI, also consistent with the findings by Tessema *et al*. (2019) (Figure 4). Interestingly, Tessema *et al*. (2019) identified Rundu district hospital as having exceptionally high within-host diversity as measured by *F*_*W S*_, which was posited to be due to a large fraction of the patients having traveled or resided in Angola. We found that Rundu district hospital had the highest mean eMOI and greatest spread across observations (mean = 4.3 [95% CI: 4.18 - 4.4], IQR = 4.88). This was mainly driven by much higher mean MOI (7 [95% CI: 6.5-7.5]), and low mean within-host relatedness (.47 [95% CI: .43 - 0.51]). The combination of high MOI and relatively low within-host relatedness, translating into high population mean eMOI, suggests that samples collected here reflect a parasite population experiencing less inbreeding and more superinfection, which may be indicative of higher transmission intensity in tandem with a larger effective population size.

**Fig. 4:**
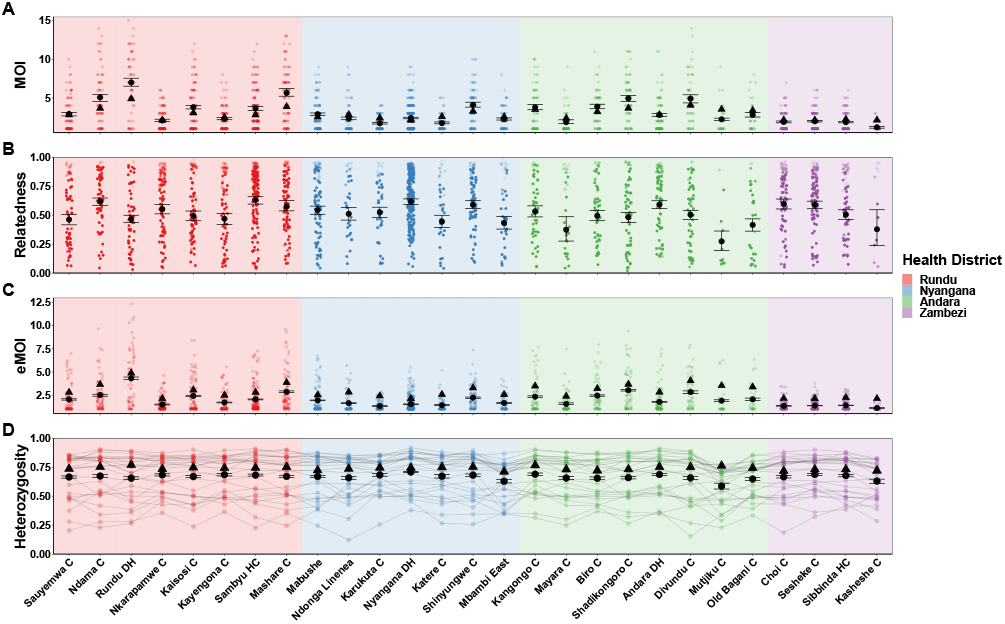
Estimated MOI, relatedness, eMOI and heterozygosity in Northern Namibia, stratified by health facility. MOIRE was run independently on data from each health facility. Two health facilities from the Zambezi region were excluded due to only having 9 samples present in each subset. Health facilities are plotted in geographic order from West to East. Plotting conventions are the same as in Figure 3.

## Discussion

Translating *Plasmodium* genetic data from naturally acquired infections into meaningful insights about population genetics or malaria transmission dynamics often begins with estimation of allele frequencies and MOI. We demonstrated through simulation that naive estimation introduces substantial biases, rendering estimation unreliable and uncomparable between settings. In particular, naive estimation systematically overestimates measures of allelic diversity such as heterozygosity and systematically underestimates MOI. State-of-the-art methods previously available to more accurately estimate individual level MOI and population allele frequencies only allow for SNP based data, and fail to directly consider within-host relatedness as an important biologic factor (Chang *et al*., 2017; Paschalidis *et al*., 2023; Ju *et al*., 2023). MOIRE fills these important gaps, demonstrating both the power and necessity of polyallelic data to obtain precise estimates of these key parameters for understanding of parasite population structure and dynamics. The R package implementing MOIRE provides a user-friendly interface for researchers to easily leverage SNP and polyallelic data to estimate these individual and population diversity metrics which are fundamental for many downstream analyses and often of direct interest themselves.

By estimating within-host relatedness, we also have introduced a new metric of diversity—eMOI—a continuous metric that integrates within-host relatedness and MOI, providing the first portable metric of within-host diversity. This metric is highly identifiable and robust to varying genetic backgrounds, and thus readily comparable across settings and genotyping technologies. We demonstrated that eMOI is a more stable metric of genetic diversity than naive MOI or *F*_*W S*_, and is insensitive to other factors that may vary across settings such as allele frequencies of given genetic markers. Further, by decomposing the genetic state of an infection into components of within-host relatedness and the number of distinct strains present, we have enabled the characterization of these quantities independently, which may be of interest in their own right. For example, within-host relatedness may be of interest in the context of understanding the role of inbreeding and co-transmission in the parasite population (Wong *et al*., 2022; Nkhoma *et al*., 2020), and the number of distinct strains may be of interest in the context of understanding superinfection dynamics.

While we have demonstrated the utility of polyallelic data, MOIRE is still compatible with SNP based data and can offer benefits over other approaches. When using SNP based panels, eMOI is still well characterized up to moderate levels, and while the reduced capacity of SNPs generally results in biased estimates, the estimates recovered reflect changes in within-host relatedness yet are stable across genetic backgrounds. Thus, these data may be useful for comparing relative ordering of eMOI across settings and providing inference. In contrast, existing analytical approaches are likely to be sensitive to model misspecification by not considering within-host relatedness and varying genetic backgrounds, and may be biased in ways that are difficult to interpret and compare across settings.

We also note that while increasing the number of loci genotyped is always beneficial, the largest gains in recovering estimates of interest are through using sufficiently diverse loci. Our simulations demonstrate that, even with a modest number of very diverse loci such as our synthetic simulations using 30 loci, eMOI can be recovered with a high accuracy and precision. Marginal increases in complexity of incorporating several highly diverse loci, for example in the context of drug resistance monitoring, may be outweighed by the substantial insights obtained from jointly understanding transmission dynamics, population structure, and drug resistance through increased accuracy of estimating resistance marker allele frequency. Modern amplicon sequencing panels have been developed precisely for these contexts, combining high diversity targets with comprehensive coverage of known resistance markers (LaVerriere *et al*., 2022; Aranda-Diaz and Neubauer Vickers, 2022; Kattenberg *et al*., 2023).

MOIRE provides a powerful tool for leveraging polyallelic data to understand malaria epidemiology, and there are multiple avenues for future work to further improve inference. First, the observation model does not currently fully leverage the information in sequencing based data where the actual number of reads may be available. This may provide additional information, e.g. to inform false positive rates by considering the number of reads attributable to an allele, as well as false negative rates by considering the total number of reads at a locus which may be indicative of sample quality. Second, we currently consider only a single, well mixed, background population parameterized by allele frequencies at each locus. However, it may be the case that there are multiple distinct populations with their own allele frequencies, and that the observed data is a mixture of these populations. This may be particularly relevant in the context of malaria transmission where there may be multiple distinct populations of parasites circulating in a region. Future work may consider a mixture model over allele frequencies, where the number of populations is a priori specified or determined through data adaptive non-parametric Bayesian modeling, and thereby identify population substructure. Alternatively, a spatially explicit approach may be feasible that would model the allele frequencies as a function of geographic location, potentially enabling resolving geographic origin of parasites within observed infections. Third, MOIRE currently assumes independence of loci. In the case of locus dependence where there is some amount of linkage disequilibrium, we would expect estimates of allele frequencies and sample specific eMOI to still be consistent if there is not a systematic bias in loci towards regions of high or low within-host relatedness.

In summary, MOIRE enables the use of polyallelic data to estimate allele frequencies, MOI, and within-host relatedness, and provides a new metric of genetic diversity, the eMOI. We have demonstrated that eMOI has improved utility, interpretability, and stability across simulated transmission settings than existing metrics of within-host diversity such as *F*_*W S*_. Furthermore, we demonstrated the utility of MOIRE through simulation and reanalysis of previously collected data, and have provided an R package to enable researchers to easily leverage polyallelic data to make inferences about malaria population dynamics. MOIRE also serves as a fundamental building block for future work, as it provides a principled approach to jointly estimate allele frequencies, MOI, and within-host relatedness from polyallelic data, which can be used as a basis for more complex modeling of population dynamics. These methods may also be of utility for other pathogens where superinfection is common, such as schistosomiasis or filarial diseases (Aemero *et al*., 2015; Hedtke *et al*., 2020).

## Acknowledgments

We thank Nicholas Hathaway for generating and sharing the regional allele frequency data used in this study.

