## supplementary for "MOIRE: A software package for the estimation of allele frequencies and effective multiplicity of infection from polyallelic data"

July 31, 2024

**2 Prior distributions** **3**

**3 Practical Considerations and Real World Data** **4**

**4 Simulation Procedure** **4**

**5 Sampling and Inference** **5**

**6 Effective MOI** **6**

**7 Regional Populations** **8**

**8 Tables** **9**

**9 Figures** **10**

### 1 Modeling transmission and observation

#### 1.1 Modeling genotyping error

We assumed that each observed sample has an intrinsic rate of false positives $\epsilon_i^+$ and false negatives $\epsilon_i^-$, reflecting the varying quality of samples that are genotyped and factors that impact ability to detect alleles accurately. This rate is divided by the number of alleles possible at each locus ($L_j$), such that the probability of observing an allele given that it is not present is $\frac{\epsilon_i^+}{L_j}$, and the probability of not observing an allele given that it is present is $\frac{\epsilon_i^-}{L_j}$. The full likelihood of observing the data given the latent true genetic state and error rates for an individual sample is then given by

$$P(X_i|Y_i,\epsilon_i^+,\epsilon_i^-)=\prod_{j=1}^{l}\prod_{a=1}^{L_j}\begin{cases}\frac{\epsilon_i^+}{L_j} & \text{if } X_{ija}=1 \text{ and } Y_{ija}=0\\ 1-\frac{\epsilon_i^+}{L_j} & \text{if } X_{ija}=1 \text{ and } Y_{ija}=1\\ \frac{\epsilon_i^-}{L_j} & \text{if } X_{ija}=0 \text{ and } Y_{ija}=1\\ 1-\frac{\epsilon_i^-}{L_j} & \text{if } X_{ija}=0 \text{ and } Y_{ija}=0\end{cases} \tag{1}$$

#### 1.2 Modeling latent genetic state

We modeled the latent genetic state as a combination of two processes. For a fixed MOI, the genetic state of each strain at a given locus can be explained either as an independent draw from the background population of parasites characterized by the population allele frequencies $\boldsymbol{\pi}$, or as being related to and explained by another within host strain. We can express this as a mixture model over the possible arrangements of $\mu_i$ strains, where each strain is either a within-host related strain at that locus or derived from the background population, giving us:

$$P(Y_i|\mu_i,r_i,\boldsymbol{\pi})=\prod_{j=1}^{l}\sum_{k=0}^{\mu_i-1}P(Y_{ij}|m=k,\mu_i,\pi_j)P(m=k|r_i,\mu_i) \tag{2}$$

Here, $m$ is a variable indicating the number of strains that are related to each other within the host, constrained to at most $\mu_i-1$ related strains, requiring at least one ”reference” strain that other strains are related to. We modeled the number of related strains at a locus within a host as a binomial random variable with probability of success $r_i$, the within-host relatedness of the infection:

$$P(m=k|r_i,\mu_i)=\binom{\mu_i-1}{k}(1-r_i)^k(r_i)^{\mu_i-k-1} \tag{3}$$

For the remaining $\mu_i-k$ unrelated strains, we modeled the genetic state of each unrelated strain as independent of the genetic state of other within host strains and dependent only on the population allele frequencies $\boldsymbol{\pi}$, allowing us to express the likelihood of the genetic state of each locus in terms of draws from a multinomial distribution:

$$P(Y_{ij}|m=k,\mu_i,\pi_j)=\sum_{y^*\in\mathcal{Y}_{Y_{ij}}}\frac{(\mu_i-k)!}{y_1^*!\cdots y_{L_j}^*!}\prod_{a=1}^{L_j}\pi_{ja}^{y_a^*} \tag{4}$$

Here, $\mathcal{Y}_{Y_{ij}}$ is the set of all possible configurations of alleles at a given locus for a given individual that are compatible with the binary vector $Y_{ij}$, where compatibility is defined as having at least one allele present in each position of the vector that is also present in $Y_{ij}$. For example, if $Y_{ij}=(1,0,1,0)$ and there are 4 unrelated strains present, then $\mathcal{Y}_{Y_{ij}} = \{(1,0,3,0),(3,0,1,0),(2,0,2,0)\}$. Naively computing this term would require enumerating all possible configurations of alleles at each locus for each individual, which may be computationally challenging for large datasets with high MOIs. We instead used a more efficient method of computing the likelihood of the latent genetic state which we describe in subsection 1.3.

##### 1.3 Alternative latent genetic state formulation

While the formulation of latent genetic state as a draw from a multinomial distribution that is then masked into a binary vector indicating presence or absence is intuitive, it is a computationally inefficient approach as we must enumerate all possible arrangements that are compatible with the latent genetic state. We can instead consider an alternative but equivalent formulation. As before, we are interested in calculating the probability of the latent genetic state, $P(Y_{ij}|m = k, \mu_i, \pi_j)$. Let $O_{ij}$ be the set of indices of alleles that are observed in sample $i$ at locus $j$. For example, if $Y_{ij} = (0,1,1,0)$ then $O_{ij} = \{2,3\}$. Let $\pi(O_{ij})$ equal the sum of allele frequencies indexed by $O_{ij}$, and $n = \mu_i - k$, the number of unrelated strains present. We can then write the probability of the latent genetic state as

$$P(Y_{ij}|m = k, \mu_i, \pi_j) = P(O_{ij}|n, \pi(O_{ij}))\pi(O_{ij})^n \tag{5}$$

where $P(O_{ij}|n, \pi(O_{ij}))$ is the probability that each allele indexed by $O_{ij}$ is present at least once after k draws, conditional on that all draws are from the set of alleles indexed by $O_{ij}$, and $\pi(O_{ij})^n$ is the probability that all $n$ alleles come from the set of alleles indexed by $O_{ij}$. The probability of every allele being present at least once is equal to 1 minus the probability that any allele is not present, which can be calculated via the inclusion exclusion principle:

$$P(O_{ij}|n, \pi(O_{ij})) = 1 - \sum_{S \subseteq O_{ij}} (-1)^{|S|-1}(1 - \sum_{a \in S} \frac{\pi_{ja}}{\pi(O_{ij})})^n \tag{6}$$

where $S$ is a subset of $O_{ij}$ and $|S|$ is the cardinality of $S$. This formulation is computationally more efficient, though less intuitive, by allowing us to calculate the probability of the latent genetic state without directly enumerating all possible arrangements of alleles that are compatible with the latent genetic state.

#### 2 Prior distributions

We assumed that sample MOIs were drawn from a zero-truncated Poisson (ZTP) distribution with rate $\lambda$.

In general, we do not know the population mean MOI $\lambda$, so we used a hyperprior distribution to model our uncertainty about $\lambda$. We assumed a gamma distribution hyperprior on $\lambda$ with user specified shape $\alpha_\lambda$ and rate $\beta_\lambda$.

Within-host relatedness was bounded between 0 and 1 and modeled with a beta distribution prior with user specified shape parameters $\alpha_r$ and $\beta_r$.

Sample specific false positive and false negative rates were bounded between 0 and user specified $\epsilon^+_{max}$ or $\epsilon^-_{max}$, respectively, and were modeled with scaled beta distributions with user specified shape parameters $(\alpha_{\epsilon^+}, \beta_{\epsilon^+})$ and $(\alpha_{\epsilon^-}, \beta_{\epsilon^-})$. We placed flat Dirichlet priors with $\alpha = 1$ on allele frequencies $\pi_j$ for each locus $j$.

### 3 Practical Considerations and Real World Data

While MOIRE is designed to incorporate the potential uncertainty from false negatives and false positives, in extreme cases there are limitations in the identifiability of their contribution separate from the presence of within-host relatedness. As within-host relatedness manifests as reduced expected diversity for a given MOI, elevated false negative rates due to strain dropout will be difficult to resolve separate from truly reduced diversity. To demonstrate this limitation, we took the whole genome sequencing data from the 3 lab constructed multi-strain mixtures in varying relative concentrations in the Pf7 dataset and extracted observed alleles corresponding to the **MaD$^4$HatTeR** genotyping panel. Because MOIRE relies on estimates of population allele frequencies to infer the latent MOI, we embedded the lab control mixture data into a larger synthetic dataset composed of 100 simulated infections with 273 highly diverse loci. We then ran MOIRE and calculated summaries of individual level parameters for the lab control mixtures (Supplementary Figure 5).

In this context of lab generated mixture controls, the concept of relatedness is not well-defined as the infections do not arise from anything resembling the assumed model of transmission. As such, the computed value of within-host relatedness is an indication of the population structure necessary for the observed data *had the infection arisen from a natural process well described by our model*, not a measure of the true within-host relatedness of the infection, which would arguably be 0 in this case as these lab strains do not share recent ancestry. So while the actual values of within-host relatedness, MOI, and subsequently eMOI, may not correspond to anything necessarily biologically meaningful in the context of these lab mixtures, they still allow us to explore potential issues with the forms of data that MOIRE is designed to handle by examining the consistency of the estimates of these quantities across the lab control mixtures.

Supplementary Figure 5 A shows the distribution of allele counts within each sample, demonstrating the shift in diversity as composition of the mixture changes due to strain dropout and alleles go undetected, so-called false negatives. More specifically, compositions that are more evenly distributed are less likely to experience allelic dropout. How this impacts the estimates of MOI and relatedness is dependent on the extent of dropout. In the case of extensive dropout, such as in the lab controls where the minor strain relative concentration is less than 10% of the total, MOIRE is unable to recover the true MOI, relatedness or eMOI (Supplementary Figure 5 B, C, D). When dropout is less extensive, MOI is relatively consistent, however the estimates of eMOI and relatedness will vary slightly depending on the extent of dropout, demonstrating potential identifiability issues of within-host relatedness when alleles are undetected.

The data presented here from the Pf7 dataset reflect whole genome sequencing data, which may be particularly sensitive to this issue as the sequencing depth at each locus is typically low, leading to higher rates of allelic dropout relative to other available technologies such as targeted amplicon sequencing. This underscores the importance of not only considering the potential for false positives and false negatives with respect to the bioinformatics pipeline, but also the sensitivity of the assay used to generate the data and its ability to detect minority strains with sufficient diversity and avoiding unusual targets such as those with copy number variation that might introduce false positives.

### 4 Simulation Procedure

We sampled population allele frequency vectors for each locus from a uniform Dirichlet distribution, or used empirical estimates from regional populations, resulting in an $L_j$-dimensional simplex for each genetic locus. For each individual, we sampled the MOI from a zero truncated Poisson (ZTP) distribution with rate $\lambda$, and then independently sampled alleles for each locus from a multinomial distribution parameterized by $p_j$. We simulated within-host relatedness for individual $i$ by sampling alleles from a randomly chosen existing strain within the host with probability $r_i$, or from the population allele frequency distribution otherwise. Relatedness $r_i$ was drawn from a beta distribution with $\alpha = 1$ and $\beta = \{.2, .6, 2\}$ for low, moderate, and highly related populations respectively. We then generated observed genetic data by applying a noisy observation process where alleles were false positives or false negatives consistent with our error model. We compared our approach to estimating allele frequencies, MOI, and observed heterozygosity to naive empirical estimators that do not consider within-host relatedness, MOI, or genotyping errors. We also evaluated the ability of our model to recover true relatedness and eMOI of samples.

### 5 Sampling and Inference

#### 5.1 Error Rates

Sample specific false positive and false negative rates were randomly initialized to a value between 0 and 0.1. False positive and false negative rates were constrained to be between 0 and 2, so sampling was conducted after transforming to an unconstrained space to improve efficiency. Proposals were sampled from a normal distribution centered around the transformed current value with a standard deviation that was adapted during burnin to achieve an acceptance rate of 0.23 with proposals accepted according to a Metropolis-Hastings acceptance probability (Gelman *et al.*, 1997; Chib and Greenberg, 1995). Priors were uniform between 0 and 1.

#### 5.2 Within-host Relatedness

Within-host relatedness was randomly initialized uniformly between 0 and 1 for each sample and constrained to be between 0 and 1, so sampling was conducted after transforming to an unconstrained space to improve efficiency. Proposals were sampled from a normal distribution centered around the transformed current value with a standard deviation that was adapted during burnin to achieve an acceptance rate of 0.23 with proposals accepted according to a Metropolis-Hastings acceptance probability. Priors were uniform between 0 and 1, reflecting our lack of prior knowledge about within-host relatedness.

#### 5.3 MOI and Latent Genotypes

Sample specific MOIs were randomly initialized to a value between the maximum number of observed alleles $\pm 3$ without exceeding specified constraints. Sample specific MOIs were constrained to be between 0 and 40. Latent genotypes were initialized to the observed genotype. Proposals for sample MOIs were generated by sampling from a symmetric distribution centered around the current value. Proposals that exceeded the constraints were rejected. Simultaneously, latent genotypes for each locus were sampled by randomly sampling the number of false positives and false negatives present, while ensuring at least one allele being a true positive, and the total number of true positives not exceeding the proposed MOI. The number of false positives and false negatives were sampled from a binomial distribution with the number of trials selected to satisfy the previously described constraints, and the probability of success equal to the false positive and false negative rates at that step, respectively. The alleles that were false positives or false negatives were randomly selected from the observed genotype. Proposals were accepted according to a Metropolis-Hastings acceptance probability.

##### 5.4 Allele Frequencies

Allele frequencies were randomly initialized on the unit simplex. Allele frequencies were then sampled according to the self adjusting logit transform proposal, also known as the SALT sampler (Director *et al.*, 2017) and accepted according to a Metropolis-Hastings acceptance probability.

##### 5.5 Mean MOI

The mean MOI was randomly initialized to a random draw from the prior distribution. The mean MOI was constrained to be between 0 and 40, so sampling was conducted after transforming to an unconstrained space to improve efficiency. Proposals were sampled from a normal distribution centered around the transformed current value with a standard deviation that was adapted during burnin to achieve an acceptance rate of 0.23 with proposals accepted according to a Metropolis-Hastings acceptance probability. The hyperprior on the mean MOI was a gamma distribution with shape and scale parameters of 0.1 and 10 respectively, reflecting our assumptions around low mean MOI.

##### 5.6 Metropolis Coupled Markov Chain Monte Carlo

It may be the case that the posterior distribution is multimodal, and that the Markov chain may get stuck in a local maxima, leading to poor mixing. To address this, we use Metropolis Coupled Markov Chain Monte Carlo ($MC^3$), also sometimes referred to as parallel tempering, or replica exchange MCMC sampling (Earl and Deem, 2005). We provide a fully parallelized implementation, allowing for the full utilization of modern high performance computing clusters or multicore desktop machines. We also leverage a non-reversible algorithm with an adaptive temperature gradient as described by (Syed *et al.*, 2022) to further improve mixing and reduce tuning required by the user.

#### 6 Effective MOI

Consider an infection with $n$ distinct unrelated strains drawn from the background population. Let $X$ be the observed genotype at a single locus with $L$ possible alleles and population frequencies $\pi$, where every allele has a non-zero frequency in the population, and $D$ be the number of distinct alleles observed at the locus

$$D = \sum_{i=1}^{L} \mathbb{I}(X_i = 1) \tag{7}$$

where $X_i = 1$ if allele $i$ is observed. By linearity of expectation, the expected number of distinct alleles (EDA) is

$$\begin{aligned}
\mathbb{E}[D] &= \sum_{i=1}^{L} \mathbb{E}[\mathbb{I}(X_i = 1)] && (8)\\
&= \sum_{i=1}^{L} \mathbb{P}(X_i = 1) && (9)\\
&= \sum_{i=1}^{L} 1 - (1 - \pi_i)^n && (10)\\
& && (11)
\end{aligned}$$

where $(1-\pi_i)^n$ is the probability that allele $i$ is not observed in any of the $n$ strains.

We note that the EDA is a metric that is dependent on MOI and allele frequencies, and is itself an interesting quantity. In the special case of $n = 2$, it is equivalent to the heterozygosity of the locus after subtracting 1. When $n > 2$, the EDA is no longer necessarily equivalent. Unsurprisingly, the EDA for a fixed n and parameterized by allele frequencies is maximized under the same conditions as the heterozygosity (when all alleles are equally likely to be observed). However, loci with differing allele frequencies (in distribution and cardinality) may have the same heterozygosity but different EDAs. This suggests that heterozygosity may be an imperfect metric of diversity in the context of mixed infections. For a fixed MOI, a higher EDA indicates greater information capacity in the locus, and thus higher precision as a tool for estimating parameters. A locus with higher EDA should be preferred when considering loci for genotyping panel development.

We also note that the EDA of a locus may not be strictly larger than the EDA of another locus across MOI, suggesting that genetic loci are more powerful for statistical purposes under limited ranges. Because of this, EDA should be considered in the context of the population distribution of MOI. In principle, it would be possible to marginalize over an estimate of the population distribution of MOI to obtain a generalized estimate of EDA that is weighted by the probability of observing a given MOI, providing a metric that is targeted to the population of interest.

The EDA also provides a natural approach to defining effective MOI. Consider an idealized locus with a heterozygosity of 1, meaning that every allele drawn from the population is unique. Practically, this is an impossibility, however we may approximate such a locus by considering the limit as the number of alleles goes to infinity and occur with equal probability. Let $L \to \infty$ and $\pi_i = \frac{1}{L}$, then the EDA for an infection with $n$ distinct strains is

$$
\begin{aligned}
\lim_{L\to\infty} E[D] &= \lim_{L\to\infty} \sum_{i=1}^{L} 1 - (1 - \frac{1}{L})^n & (12)\\
&= \lim_{L\to\infty} L - L(\frac{L-1}{L})^n & (13)\\
&= n & (14)
\end{aligned}
$$

As expected, in the limit of an infinitely diverse locus, the EDA is equal to the MOI. We now consider the EDA for an infinitely diverse locus in an infection with within-host relatedness $r$. While MOI remains a fixed quantity, the number of related strains is now a binomial distributed random variable with parameters $n-1$ and $r$. We define the effective MOI (eMOI) as the expected number of distinct alleles observed in an infection with $n$ distinct strains and within-host relatedness $r$ at a locus with infinite diversity. By construction, only unrelated strains contribute to the EDA, thus the eMOI is simply the EDA marginalized over the distribution of unrelated strains in an infection with $n$ distinct strains and within-host relatedness $r$

$$
\begin{aligned}
\text{eMOI} &= \sum_{k=0}^{n-1} \binom{n-1}{k} r^k (1-r)^{n-1-k}(n-k) & (15)\\
&= 1 + (1-r)(n-1) & (16)
\end{aligned}
$$

As $r$ goes to zero, this quantity approaches the MOI, and as $r$ goes to one, this quantity approaches 1, recovering a natural measure of within-host diversity that is sensitive to both MOI and within-host relatedness.

### 7 Regional Populations

In order to assess potential real world performance of MOIRE, we simulated genotyping data from 12 regional populations parameterized by the MalariaGEN Pf7 dataset (?). Regional populations were defined by applying t-distributed stochastic neighbor embedding (t-SNE) to the full Pf7 dataset then clustering countries into regional populations. Countries included in each regional population are summarized in Table 2. Marginal allele frequencies were naively estimated within each region for each genotyping panel from whole genome sequencing data available from the Pf7 dataset. We have made these allele frequency estimates available as supplementary data in our R package, accessible as `moire::regional_allele_frequencies`. Alleles within regions with a frequency below 1% were excluded from microhaplotype panels to avoid excessive computational burden when running MOIRE. Uninformative loci (only 1 allele observed) were removed for each region. Interquartile ranges of the number of alleles observed and the number of loci in each panel are summarized in Table 3. Summaries of panel diversity, evaluated by calculating the expected number of distinct alleles from 2 to 10 possible strains, are provided in figure 4. We note that not all unique alleles included in the simulation may be reliably genotypable, e.g. homopolymers and tandem repeats, depending on the fidelity of amplification and sequencing. Simulations may provide an optimistic estimate of performance.

### 8 Tables

| **Panel** | Mean MOI (Cov. %) | Mean Relatedness (Cov. %) | Mean eMOI (Cov. %) |
|---|---|---|---|
| **100 SNP** | 1.18 (.17) | 0.24 (.05) | 0.26 (.23) |
| **Moderate Div.** | 0.67 (.24) | 0.15 (.06) | 0.07 (.49) |
| **High Div.** | 0.38 (.29) | 0.10 (.09) | 0.04 (.49) |
| **Very High Div.** | 0.29 (.27) | 0.08 (.11) | 0.04 (.34) |
| **24 SNP** | 0.63 (.21) | 0.23 (.06) | 0.32 (.08) |
| **101 SNP** | 0.53 (.30) | 0.20 (.09) | 0.33 (.09) |
| **AMPLseq** | 0.39 (.23) | 0.11 (.15) | 0.09 (.23) |
| **MaD$^4$HatTeR** | 0.42 (.28) | 0.11 (.27) | 0.09 (.28) |
| **AmpliSeq** | 0.40 (.33) | 0.10 (.24) | 0.08 (.33) |

Table 1: Mean absolute deviation (MAD) and CI coverage at the 95% interval of estimates of population mean MOI, mean within-host relatedness, and mean eMOI across all simulations using synthetic (top) and real-world (bottom) genotyping panels.

| **Region** | Countries |
|---|---|
| South America - Central | Brazil, French Guiana, Guyana, Peru, Venezuela |
| South America - North | Colombia |
| South Asia | Bangladesh, India |
| South East Asia - East | Cambodia, Laos, Vietnam |
| South East Asia - West | Myanmar, Thailand |
| Papua New Guinea | Papua New Guinea |
| Central Africa | DRC, Zambia |
| Central West Africa | Cameroon, Gabon, Nigeria |
| West Africa | Benin, Burkina Faso, Ghana, Guinea, Ivory Coast, Mali, Mauritania, Senegal, The Gambia |
| East Africa | Ethiopia, Kenya, Madagascar, Sudan, Tanzania, Uganda |
| South Africa | Namibia |
| South East Africa | Malawi, Mozambique |

Table 2: Countries included in each regional population.

| Region | AMPLseq | MaD$^4$HatTeR | AmpliSeq |
| --- | --- | --- | --- |
| South America - Central | 2-3 (n=106) | 2-3 (n=141) | 2-4 (n=128) |
| South America - North | 2-3 (n=107) | 2-3 (n=136) | 2-3 (n=114) |
| South Asia | 3-7 (n=121) | 3-7 (n=165) | 2-9 (n=155) |
| South East Asia - East | 2-5 (n=106) | 2-5 (n=137) | 2-7 (n=129) |
| South East Asia - West | 2-5 (n=110) | 2-4 (n=147) | 2-5 (n=148) |
| Papua New Guinea | 2-4 (n=115) | 2-5 (n=151) | 2-6 (n=148) |
| Central Africa | 3-8 (n=121) | 4-7 (n=165) | 2-10 (n=158) |
| Central West Africa | 3-8 (n=121) | 3-7 (n=164) | 2-10 (n=155) |
| West Africa | 2-8 (n=122) | 3-7 (n=164) | 2-11 (n=149) |
| East Africa | 3-8 (n=122) | 4-8 (n=165) | 2-11 (n=156) |
| South Africa | 3-9 (n=125) | 4-8 (n=165) | 2-7 (n=161) |
| South East Africa | 3-8 (n=119) | 4-7 (n=165) | 2-11 (n=153) |

Table 3: Interquartile range (IQR) of the number of alleles per locus for each region and genotyping panel. Number of loci included in each panel is indicated in parentheses.

### 9 Figures

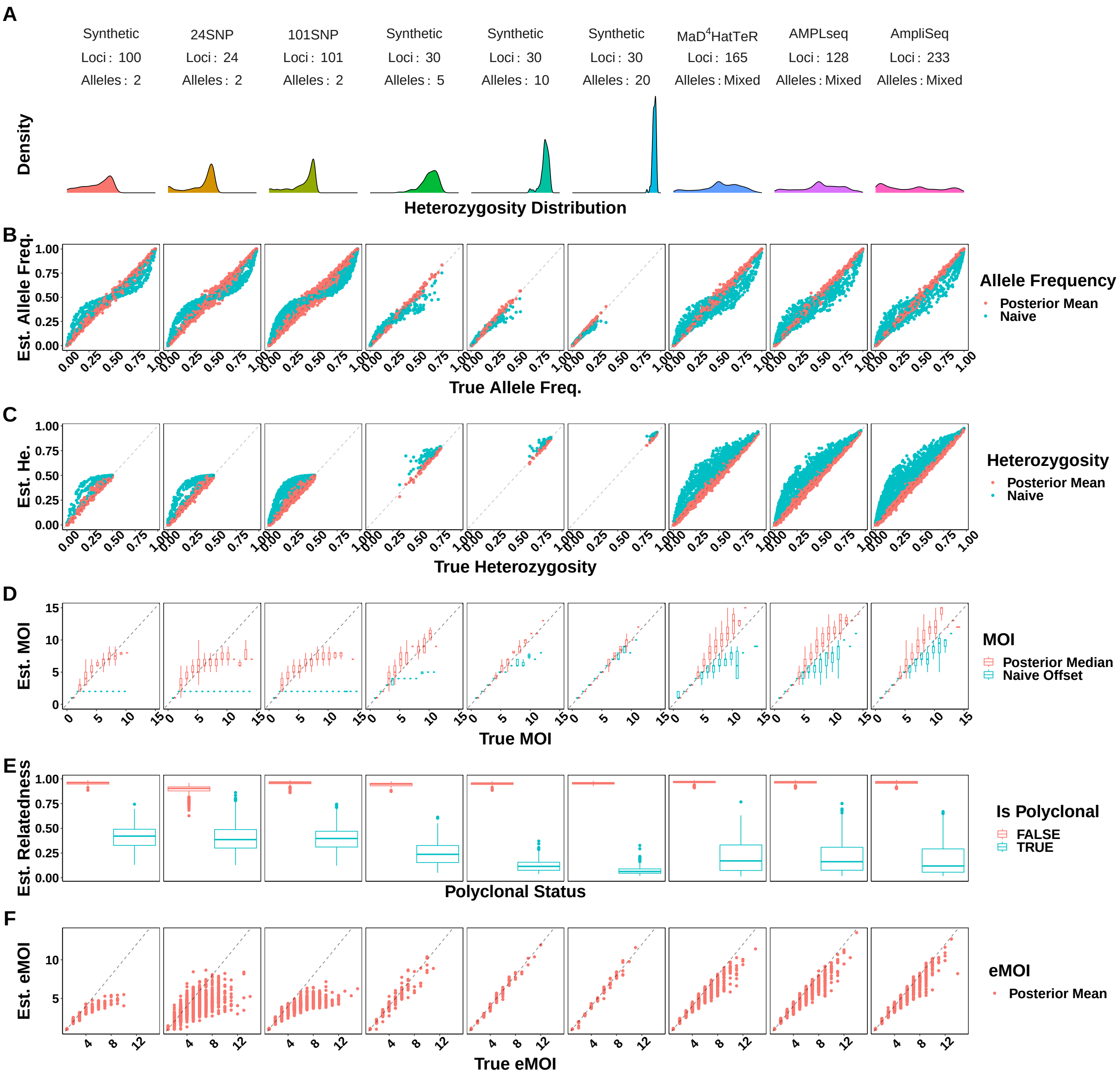

Figure 1: **Attempting to estimate relatedness in the absence of relatedness did not introduce bias into parameters of interest such as MOI, heterozygosity, and allele frequencies.** Simulations were pooled across mean MOIs. False positive and false negatives rates were fixed to 0.01 and 0.1 respectively.

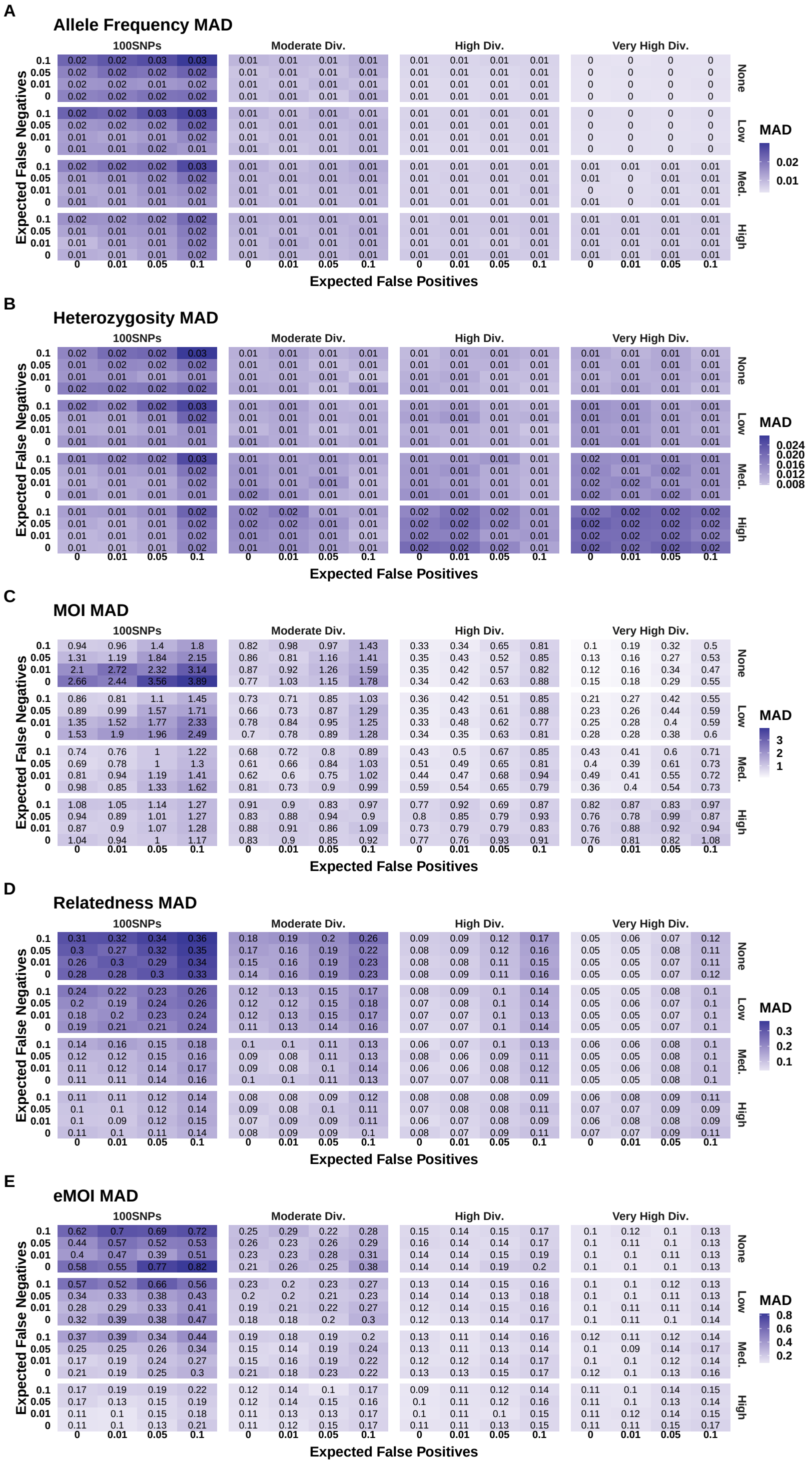

Figure 2: **Mean absolute deviation (MAD) of parameter estimates by MOIRE across panels of varying genetic diversity and stratified by population levels of within-host relatedness.**

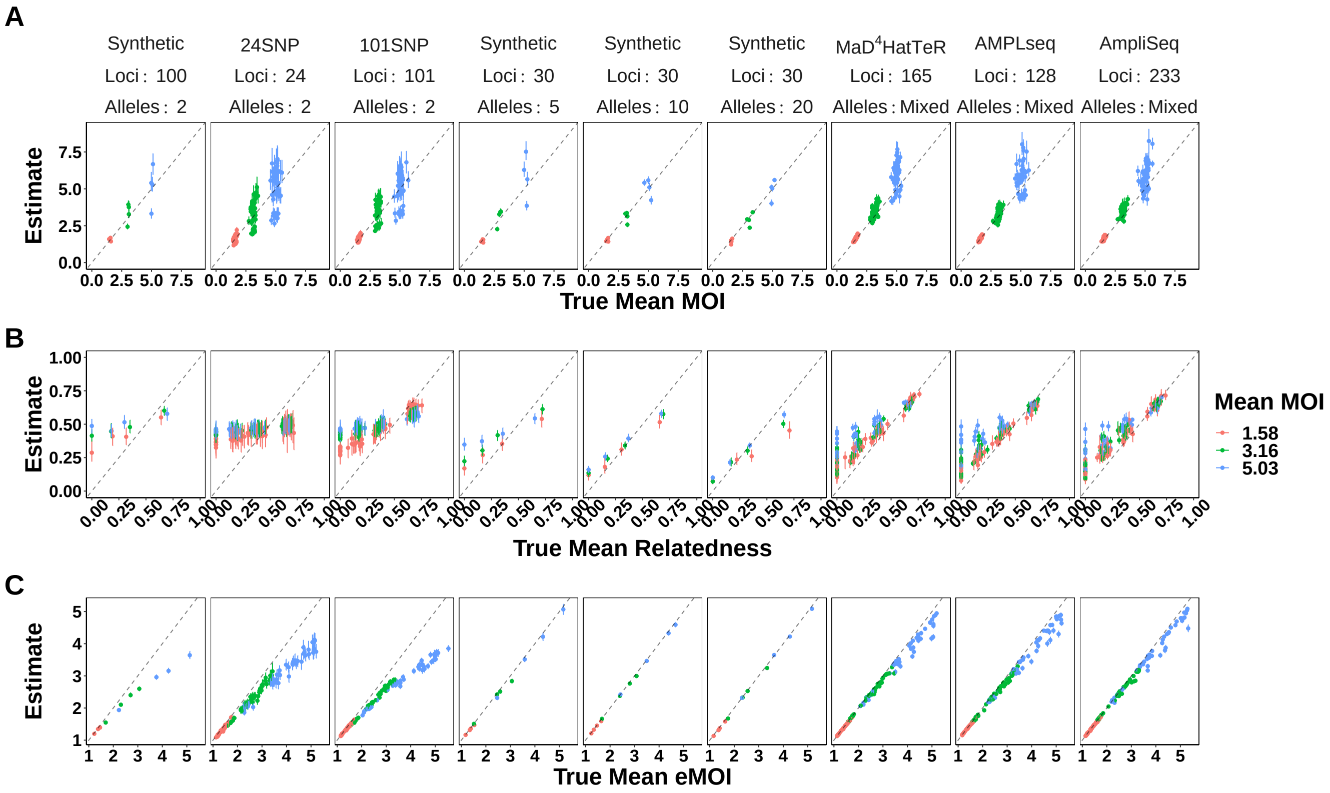

Figure 3: **True vs. estimated values of population level parameters across panels of varying genetic diversity.** Each symbol represents the estimated value of the population summary parameter for a single simulated dataset, with the true value of the parameter on the x-axis and the estimated value on the y-axis. False positive and false negatives rates were fixed to 0.01 and 0.1 respectively. MOIRE accurately recovered population mean MOI and eMOI, as well as mean relatedness when relatively high. Overestimation of mean relatedness at low true values did not result in a significant bias in eMOI when sufficiently diverse genetic markers were used.

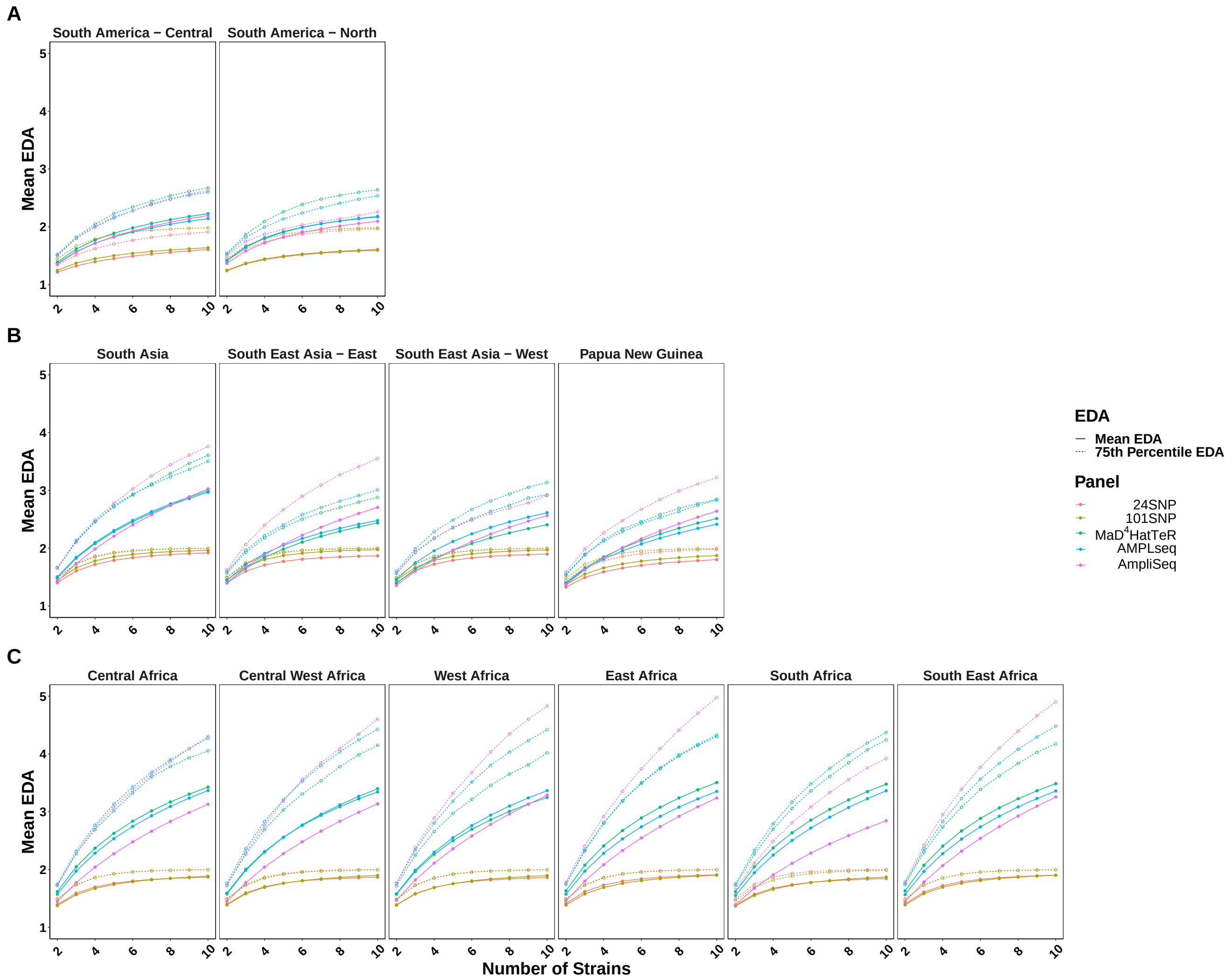

Figure 4: **Mean expected number of distinct alleles (EDA) across 12 regional populations for 5 selected genotyping panels.** Marginal allele frequencies within each regional population were estimated from the MalariaGEN Pf7 dataset. We then evaluated the expected number of distinct alleles for each locus within each regional population for each genotyping panel. The solid lines indicate the mean EDA across loci within each regional population, and the dashed lines indicate the 75$^{\text{th}}$ percentile. We note that heterozygosity for each panel is equal to the EDA minus 1 when the number of strains is 2.

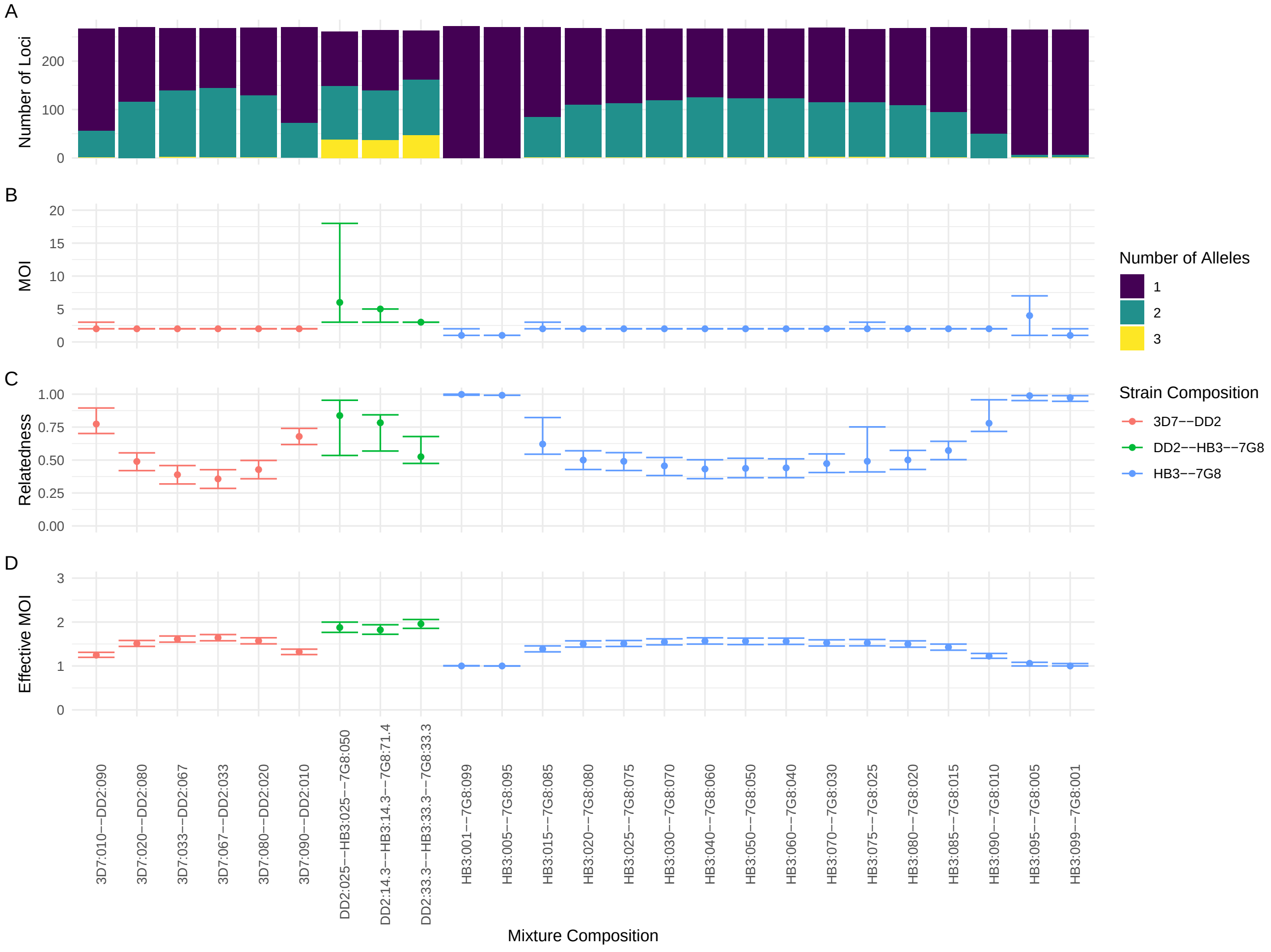

Figure 5: **Estimation of individual level parameters for lab control mixtures from the Pf7 dataset.** Figure A shows the distribution of allele counts within each sample, demonstrating the shift in diversity as composition of the mixture changes due to strain dropout. Figure B, C, and D show the posterior distribution of eMOI, MOI, and within-host relatedness for each sample respectively. Points are colored by the strain composition of the sample and indicate the median of the posterior distribution. Intervals around the points indicate the 95% credible interval. Relative frequencies of the strains are indicated within labels on the x-axis. We note that the true values of eMOI and relatedness are not known for these lab control mixtures as they are defined with respect to a background population, however we may still evaluate the consistency of the estimates of these quantities across the lab control mixtures.
